# Amphetamine alters an EEG marker of reward processing in humans and mice

**DOI:** 10.1101/2021.08.25.457689

**Authors:** James F. Cavanagh, Sarah Olguin, Jo A Talledo, Juliana E. Kotz, Benjamin Z. Roberts, John A Nungaray, Joyce Sprock, David Gregg, Savita G. Bhakta, Gregory A. Light, Neal R. Swerdlow, Jared W. Young, Jonathan L. Brigman

## Abstract

The bench-to-bedside development of pro-cognitive therapeutics for psychiatric disorders has been mired by translational failures. This is in part due to the absence of pharmacologically-sensitive cognitive biomarkers common to humans and rodents. Here, we describe a cross-species translational marker of reward processing that is sensitive to the dopamine agonist, d-amphetamine. Motivated by human electroencephalographic (EEG) findings, we recently reported that frontal midline delta-band power is also an electrophysiological biomarker of reward surprise in mice. In the current series of experiments, we determined the impact of parametric doses of d-amphetamine on this reward-related EEG response from humans (n=23) and mice (n=28) performing a probabilistic learning task. In humans, d-amphetamine (placebo, 10 mg, 20 mg) boosted the Reward Positivity event-related potential (ERP) component as well as the spectral delta-band representations of this signal. In mice, d-amphetamine (placebo, 0.1 mg/kg, 0.3 mg/kg, 1.0 mg/kg) boosted both reward and punishment ERP features, yet there was no modulation of spectral activities. In sum, the present results confirm the role of dopamine in the generation of the Reward Positivity in humans, and paves the way towards a pharmacologically valid biomarker of reward sensitivity across species.

## Introduction

Animal models of cognitive processes have been beset by translational difficulties. We recently addressed this problem by identifying common electrophysiological biomarkers of isolated cognitive processes in both humans and mice (Cavanagh et al. 2021). One of the most promising outcomes identified in that project was a common delta-band spectral reflection of the *Reward Positivity* (RewP), an event-related potential (ERP) component that has been well-described in human participants (Holroyd et al. 2008; Proudfit 2015). While the discovery of cross-species biomarkers is an important step, their translational utility requires a demonstration of consistent pharmacological predictive validity (i.e. similar drug effects across species). Here, we tested whether this cortical reward signal had similar sensitivity to d-amphetamine in mice and humans.

The RewP is a positive deflection in the ERP that is most commonly quantified over fronto-central sites around 200-400ms after reward presentation (Holroyd et al. 2011; Proudfit 2015; Heydari and Holroyd 2016). Both the RewP and its delta-band spectral reflection scale with the degree of positive reward prediction error, whereby “better than expected” outcomes evoke increasingly larger RewP amplitudes (Baker and Holroyd 2011; Cavanagh 2015; Holroyd and Umemoto 2016). These two criteria are notable: they indicate that this reward-related signal is specific and sensitive to positive reward prediction error, fulfilling the stringent criteria of being an invariant neural marker of this computational process (Cacioppo and Tassinary 1990).

The RewP is diminished in disorders like major depression (Bress et al. 2013; Webb et al. 2016) and Parkinson’s disease (Brown et al. 2020b), suggesting it might be a trans-diagnostic biomarker of reward responsiveness, learning, and valuation. The emerging mechanistic understanding of this signal increases its appeal: the reward prediction error computation reflected by the RewP mirrors the dopamine-driven learning process underlying reinforcement learning (Sutton and Barto 1998). The cortically-generated RewP has been theorized to be modulated by phasic midbrain dopamine (Holroyd et al. 2008), however empirical tests of this have been sparse and beset by methodological complications (see discussion), with little evidence for pharmacological predictive validity across species. It remains unknown if these cortical and midbrain systems influence each other in a causal manner, if they are both influenced by a third variable, or if they reflect parallel processes. Despite these uncertainties, the current understanding of the RewP suggests that it should be affected by manipulations of dopaminergic activity. The current study aimed to provide an initial test of this intuitive hypothesis, as well as facilitate the first test of common underlying mechanisms in the homologous mouse cortical signal.

## Materials and Methods

### Human Participants

Healthy women and men between the ages 18-35 years were recruited from the community via public advertisements and compensated monetarily for study participation. This study was conducted at the UCSD Medical Center and was approved by the UCSD Human Subject Institutional Review Board. Participants first completed a phone screen to assess current and past medical and psychiatric history, medication and recreational drug use and family history of psychosis. During the subsequent in-person screening visit, participants signed the consent form and completed assessments of physical and mental health, including a structured clinical interview, self-report questionnaires about caffeine intake and handedness, a hearing test, physical examination, an electrocardiogram, urine toxicology screen, urine pregnancy test for females, the MATRICS Comprehensive Cognitive Battery (MCCB) and a Wide Range Achievement Test (WRAT) for IQ assessment. See Supplemental Table 1 for these clinical demographics. A total of 12 male and 11 female participants completed all three sessions and are included here.

A double blind, randomized, placebo-controlled, counterbalanced, within-subject design was utilized. Participants received either placebo or one of two active doses of d-amphetamine (10 or 20 mg) orally on each of the three test days which were separated by one week. On all assessment days, participants arrived at 8:30 am after overnight fasting, completed a urine toxicology screen and a urine pregnancy test in females, and ate a standardized breakfast. Vital signs and subjective symptom rating scale scores were obtained at specific intervals pre- and post-administration. Starting 120 minutes post-administration, subjects completed cognitive tests and simultaneous EEG recording. See Supplemental Table 2 for experimental session information.

### Probabilistic Learning Task (PLT)

This version of the PLT was identical to our previous investigation (Cavanagh et al. 2021), see Figure 1A. Participants were presented a stimulus pair (e.g., bicycle/phone, chair/clip, plug/flashlight) on a computer monitor and instructed to select the “target” stimulus using a digital 4-switch USB arcade-style joystick. Participants were given feedback after each trial about whether their response was “correct” or “incorrect”. Reward probabilities for the target / non-target stimulus were set within a block of 60 trials (80/20, 70/30, 60/40, 50/50), but stimuli differed between trial blocks (first block was bicycle/phone at 80/20, then the next block was chair/clip at 60/40, etc.). Overall performance was calculated as the total number of correct target selections in each block of 60 trials.

**Figure 1.**
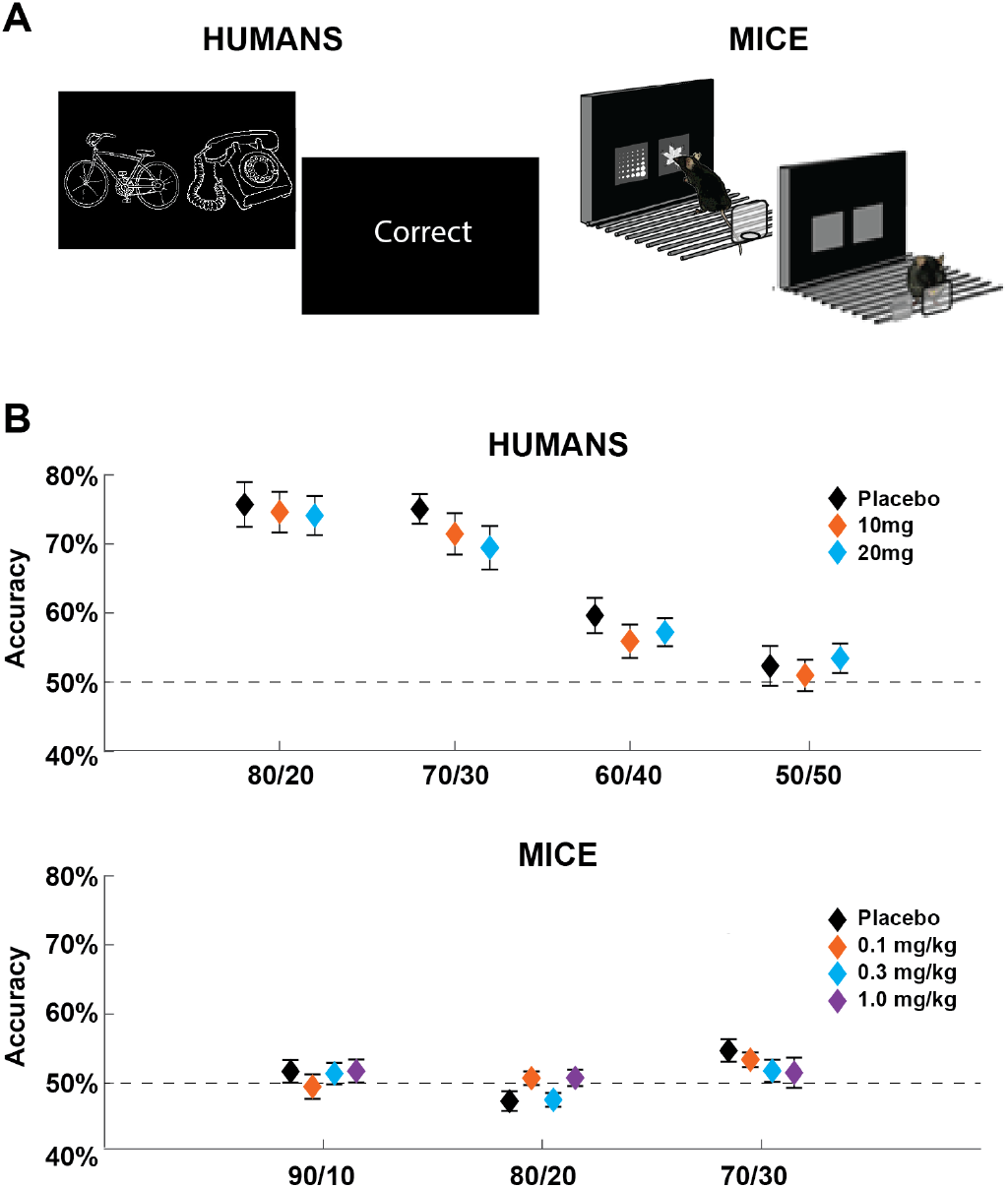
Task and performance. A) The Probabilistic Learning Task required the subject to select the stimulus that probabilistically led to reward most often. In humans and mice, each trial required a choice between two stimulus icons. B) Performance, split by block and drug. Humans displayed approximate matching behavior with no difference in performance due to amphetamine dose. In the 50/50 condition there was no chance to learn a ‘better’ stimulus and performance was at chance. Mice struggled to reliably discriminate between each pair of stimuli, but they did perform better than chance in aggregate.

### Human Electrophysiological Recording and Pre-Processing

Continuous electrophysiological (EEG) data were recorded in DC mode from 64 scalp leads using a BioSemi Active Two system. Four electrooculograms (EOG) recorded at the superior and inferior orbit of the left eye and outer canthi of each eye, and one nose and two mastoid electrodes were also collected. All data were collected using a 1048 Hz sampling rate utilizing a first-order anti-aliasing filter. Custom Matlab scripts and EEGLab (Delorme and Makeig 2004) functions were used for all data processing. Data were first down sampled to 500 Hz, epoched around the imperative cues (−2000 to +5000 ms) and then average referenced. Bad channels and bad epochs were identified and were subsequently interpolated and rejected respectively, then blinks were removed following independent component analysis.

### Animal Subjects

Female and male C57BL/6J mice were obtained from The Jackson Laboratory (Bar Harbor, ME), housed in same sex groupings of 2 per cage in a temperature- and humidity-controlled vivarium under a reverse 12 h light/dark cycle (lights off 0800 h) and tested during the dark phase. All experimental procedures were performed in accordance with the National Institutes of Health Guide for Care and Use of Laboratory Animals and were approved by the University of New Mexico Health Sciences Center Institutional Animal Care and Use Committee. A total of 15 male and 13 female mice were used. See Supplemental Methods for information on touchscreen pre-training and surgery.

A randomized, placebo-controlled, counterbalanced, within-subject design was utilized. Mice received either placebo or one of three active doses of d-amphetamine (0.1 mg/kg, 0.3 mg/kg, 1.0 mg/kg) on each of the four test days which were separated by a 72 hours washout period. Starting 15 minutes post-administration, subjects completed the PLT with simultaneous EEG recording. Notably, since 1.0 mg/kg leads to significantly increased hyperactivity and significantly decreased task focus, it was investigated as an upper limit of dosing.

### Mouse PLT

This version of the PLT was also identical to our previous investigation (Cavanagh et al. 2021), see Figure 1A. Each session, mice were presented with three pairs of unique stimuli (fan/marble, spider/fan, honey/cave) in three separate 20 trial blocks, for a total for 60 trials per session. For the first block, the target stimulus was rewarded 90% of the time and the non-target stimulus was rewarded 10% of the time. The next blocks included 80/20 and then 70/30 reinforcement rates. The mice were given 120 minutes to complete the task. Rewards consisted of the immediate delivery of an auditory tone for 1 second signaling the availability of liquid reward (30μl strawberry Nesquik, Nestle). Punishments consisted of immediate illumination of the house light for 10 seconds before the next trial could be initiated.

### Human and Mouse EEG Processing

For the sake of descriptive simplicity, both the scalp-recorded signal in humans as well as the dura-recorded signal in mice are referred to as ‘EEG’. Time-Frequency measures were computed by multiplying the fast Fourier transformed (FFT) power spectrum of single trial EEG data with the FFT power spectrum of a set of complex Morlet wavelets defined as a Gaussian-windowed complex sine wave: e^i2πtf^e^-t^2/(2xσ^2)^, where t is time, f is frequency (which increased from 1-50Hz in 50 logarithmically spaced steps) and the width (or ‘cycles’) of each frequency band were set to increase from 3/(2πf) to 10/(2πf) as frequency increased. Then, the time series was recovered by computing the inverse FFT. The end result of this process is identical to time-domain signal convolution, and it resulted in estimates of instantaneous power taken from the magnitude of the analytic signal. Each epoch was then cut in length from −500 to +1000 ms peri-feedback.

Averaged power was normalized by conversion to a decibel (dB) scale (10*log10[power(t)/power(baseline)]), allowing a direct comparison of effects across frequency bands. The baseline consisted of averaged power −300 to −200 ms before all imperative cues. A 100 ms duration is often used as an effective baseline in spectral decomposition since pixel-wise time-frequency data points have already been resolved over smoothed temporal and frequency dimensions with the wavelets. For ERPs, a baseline correction of −300 to −200 ms was applied and data were 20 Hz low-pass filtered prior to averaging. Feedback-locked analysis was conducted at electrode FCz in humans and the frontal lead in rodents.

### Statistical Analysis

Analytic methods are either identical to our previous report (Cavanagh et al. 2021), or they are new analyses. Analysis of the same time-frequency region of interest (tf-ROI) was proposed to be a major constraint within our ‘learn-confirm’ strategy between experimental phases (Cavanagh et al. 2021). ERPs were not reported in our prior ‘learn phase’ investigation; we now show the relevant mouse ERPs from that previous investigation in Supplemental Figure 1. We also report additional analyses that were facilitated by this novel ‘confirm’-stage multi-session, within subject design. As in our previous report, hypotheses were specific to the reward-related EEG; however, punishment-related EEG signals are again reported here for consistency. The number of epochs for each condition are shown in Supplemental Figure 2.

Species were analyzed with separate mixed effects models. Each mouse was treated as a random effect, similar to humans. Drug condition and sex were treated as fixed effects, although there were no *a priori* hypotheses for sex effects. Given the established effects of dopamine agonism on frontal cortical activities, either linear or quadratic effects could be expected (depending on dose and individual differences). Fitting with the demands of parsimony and the inferential affordance allowed by the number of drug conditions, this resulted in linear contrasts for humans (placebo, 10 mg, 20 mg) but quadratic contrasts for mice (placebo, 0.1 mg/kg, 0.3 mg/kg, 1.0 mg/kg).

Scalar values of spectral power were derived from the same spatial, temporal, and frequency windows as in our foundational paper. For these expectation-based contrasts, comparisons were split based on the probabilistic aspect of the reward feedback, creating high probability (i.e. target response followed by reward) vs. low probability (i.e. non-target response followed by reward) contrasts. We adhered to our previously-defined tf-ROIs and we again omitted data from the 50/50 condition in humans (human reward: 1.3 Hz to 2 Hz, 250 to 550 ms; human punishment: 4 to 6.5 Hz, 450 to 750 ms; mouse reward: 1 to 4 Hz, 250 to 550 ms; mouse punishment: 4.5 to 7.5 Hz, 500 to 800 ms). However, this study design offered an additional chance for inference.

Unlike our previous study, this multi-session design facilitated a comparison of similar feedback processes across levels of the fixed effect (drug) without the requirement of deriving a performance-based contrast to serve as a fixed contrast (expectation). This design enabled a comparison of the grand average feedback-related response outside the constraints of behavioral performance. This is beneficial, given the poor performance of mice on the task (Fig 1B). Critically, human feedback-locked EEG activities are most often assessed in un-learnable tasks with 50/50 reward probabilities (e.g. the two-door task: Foti and Hajcak 2009; Angus et al. 2015; Proudfit 2015; Mulligan and Hajcak 2017), highlighting how learning is not a necessary requirement for eliciting these cortical signatures of feedback receipt.

Due to this affordance, here we investigated grand average ERPs for the first time. ERP components were *de novo* defined based on the grand average over all drug conditions (human reward: 250 to 400 ms, human punishment: 250 to 500 ms. mouse feedback: 400 to 600 ms), see Figure 2. In humans the grand average time-frequency activity peaked earlier than the expectation-defined contrasts, so we defined new time ranges based on the grand average over all drug conditions. For reward, this earlier window was from 100 to 400 ms and for punishment this earlier window was 250 to 550 ms. There was no apparent simple grand average peak for feedback in mice, so the previously defined tf-ROIs were used in all analyses.

**Figure 2.**
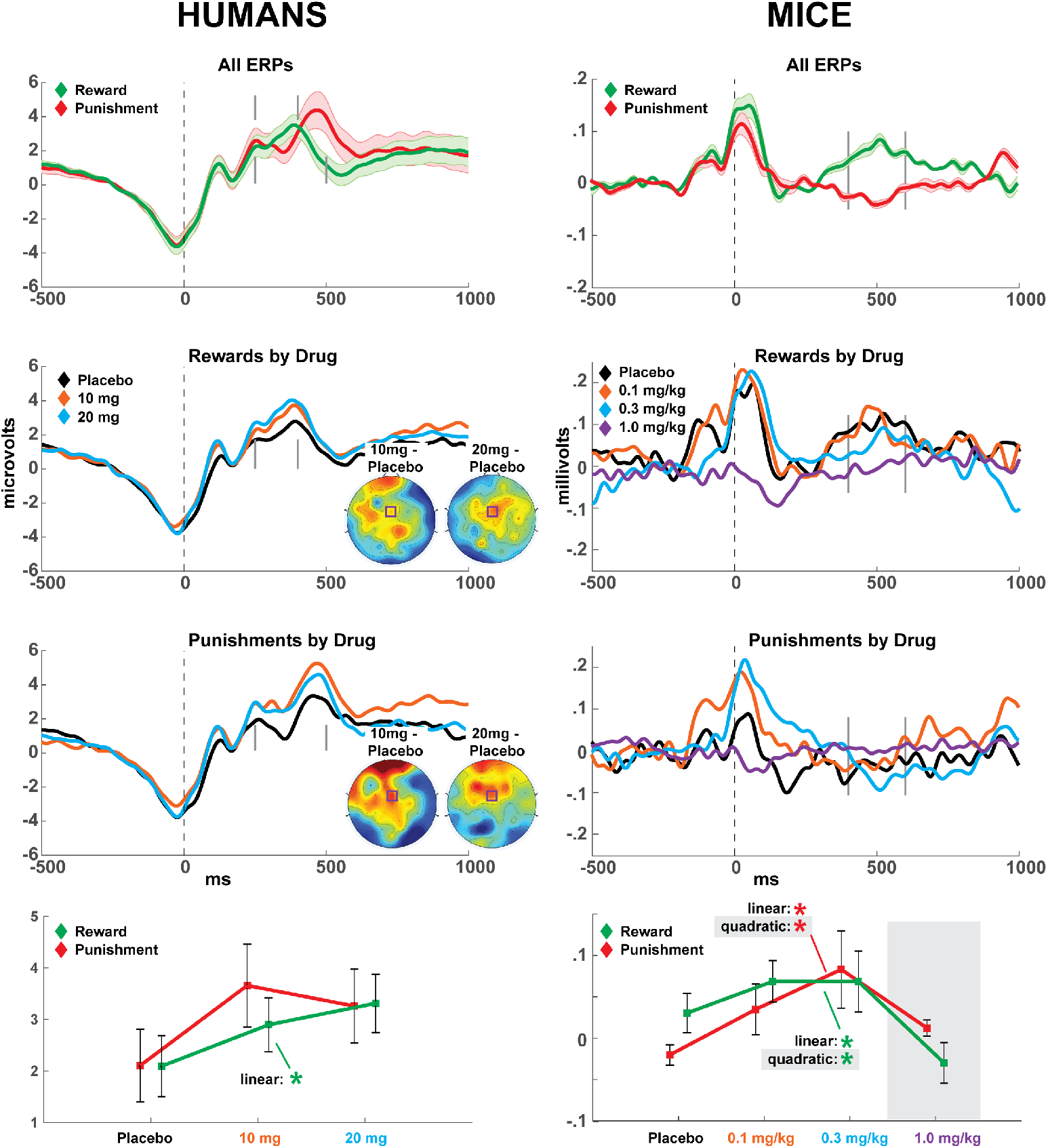
Feedback-locked ERPs from humans and mice. Grand average plots show feedback-locked ERP activity averaged over all drug conditions. Time windows for quantification are show with grey bars. Below, each feedback condition ERP is shown, split by drug dose. The same time window quantifications are shown with grey bars. Human ERPs topographical plots demonstrate drug condition differences; the purple square identifies the FCz electrode that was *a priori* chosen for analysis. Line plots (mean +/- SEM) detail the drug effects on each ERP component. In humans, only the RewP component was significantly affected by d-amphetamine. Mouse ERP features were characterized by quadratic effects with increasing d-amphetamine dose, however once the ceiling-level condition of 1.0 mg/kg was removed a similar linear trend as humans was also statistically identified. *p<.05 **p<.01.

## Results

### Performance

The full statistical results for performance data are reported in Supplemental Table 3. In *humans*, there was a main effect of block (easier pairs were more accurate, see Figure 1B) and a drug*sex interaction, which revealed a tendency for females to perform slightly worse under higher amounts of d-amphetamine whereas males were unaffected (see Supplemental Fig 3). In *mice*, there was no effect of block nor any effect with drug. This indicates that mice had poor performance discriminating the probabilities between difficulty conditions, although they managed to perform above chance when all drug and block conditions were collapsed (one sample t-test vs. chance: t(27)=2.15, p=.04), see Figure 1B.

### ERPs

All statistical results for ERP data are reported in Table 1. In *humans*, there was a main effect of drug with a linear trend for increased RewP amplitude with increasing dose of d-amphetamine. This effect was not present in the corresponding punishment condition, which showed a trend towards a quadratic effect (although the quadratic contrast was not significant either). In *mice*, there was a quadratic effect of drug on reward and punishment ERP amplitudes. Since 1.0 mg/kg of d-amphetamine was at ceiling-level tolerance and is thus not ideally suitable for comparison to humans, we examined if the same linear effect observed in humans was present in the first three drug conditions alone. Similar to the pattern observed in humans, there was a linear effect of drug for the first three conditions for both reward and punishment ERP features.

**Table 1.** Mixed linear model outcomes for ERP amplitudes, grand average tf-ROI power, and expectation-contrasted (low probability vs. high probability feedback) tf-ROI power. Quadratic effects were not tested in humans. An additional column details the main linear effect for drug dose in mice when omitting the 1.0 mg/kg (ceiling) condition in mice.

|  |  | Main:<br>Drug | Main:<br>Sex | Drug*Sex | Drug*Drug | 3-Way | Drug<br>– no 1.0 mg |
| --- | --- | --- | --- | --- | --- | --- | --- |
| Human:<br>Reward | ERP | **F=4.53**<br>**df=19.67**<br>**p=.046** | F=1.91<br>df=34.12<br>p=.18 | F=.93<br>df=19.67<br>p=.35 |  |  |  |
|  | tf-ROI<br>Grand<br>Average | **F=6.83**<br>**df=22.12**<br>**p=.02** | **F=10.42**<br>**df=20.99**<br>**p=.004** | F=1.92<br>df=22.12<br>p=.18 |  |  |  |
|  | tf-ROI<br>Expectation<br>Contrast | **F=4.73**<br>**df=28.21**<br>**p=.04** | **F=8.73**<br>**df=34.33**<br>**p=.006** | **F=4.55**<br>**df=28.21**<br>**p=.04** |  |  |  |
| Human:<br>Punishment | ERP | F=3.04<br>df=18.69<br>p=.10 | F=.39<br>df=29.80<br>p=.54 | F=.49<br>df=18.69<br>p=.49 |  |  |  |
|  | tf-ROI<br>Grand<br>Average | **F=8.30**<br>**df=23.36**<br>**p=.008** | F=.60<br>df=30.26<br>p=.45 | F=2.06<br>df=23.36<br>p=.16 |  |  |  |
|  | tf-ROI<br>Expectation<br>Contrast | F=2.84<br>df=32.68<br>p=.10 | F=2.22<br>df=42.48<br>p=.14 | F=1.63<br>df=32.68<br>p=.21 |  |  |  |
| Mouse:<br>Reward | ERP | F=0.20<br>df=55.78<br>p=.66 | F=2.19<br>df=38.61<br>p=.15 | F=.09<br>df=55.78<br>p=.76 | **F=5.62**<br>**df=36.28**<br>**p=.02** | F=0.84<br>df=36.28<br>p=.37 | **F=5.58**<br>**df=22.75**<br>**p=.03** |
|  | tf-ROI<br>Grand<br>Average | F=0.73<br>df=45.70<br>p=.40 | F=1.17<br>df=60.45<br>p=.28 | F=2.88<br>df=45.70<br>p=.10 | F=1.14<br>df=72.28<br>p=.29 | F=0.14<br>df=72.28<br>p=.71 | F=0.97<br>df=25.66<br>p=.33 |
|  | tf-ROI<br>Expectation<br>Contrast | F=0.57<br>df=75.05<br>p=.46 | F=0.36<br>df=60.93<br>p=.55 | F=0.34<br>df=75.05<br>p=.56 | F=1.73<br>df=102.08<br>p=.19 | F=0.80<br>df=102.08<br>p=.37 | F=.06<br>df=53.92<br>p=.81 |
| Mouse:<br>Punishment | ERP | F=3.81<br>df=51.22<br>p=.06 | F=.33<br>df=48.16<br>p=.57 | F=.47<br>df=51.22<br>p=.50 | **F=4.19**<br>**df=57.23**<br>**p=.045** | F=0.76<br>df=57.23<br>p=.39 | **F=6.14**<br>**df=46.19**<br>**p=.02** |
|  | tf-ROI<br>Grand<br>Average | F=0.01<br>df=76.93<br>p=.95 | F=1.55<br>df=52.77<br>p=.21 | F=0.001<br>df=76.93<br>p=.97 | F=0.00<br>df=65.48<br>p=.99 | F=0.28<br>df=65.48<br>p=.60 | F=0.04<br>df=38.24<br>p=.84 |
|  | tf-ROI<br>Expectation<br>Contrast | F=0.95<br>df=64.28<br>p=.33 | F=0.05<br>df=59.78<br>p=.82 | F=0.05<br>df=64.28<br>p=.82 | F=1.15<br>df=70.07<br>p=.29 | F=1.00<br>df=70.07<br>p=.32 | F=4.03<br>df=37.19<br>p=.052 |

### TF-ROI: Grand Average

All statistical results for grand average tf-ROIs are reported in Table 1. In *humans*, there were main effects of drug with linear trends for increased reward delta power and punishment theta power with increasing dose of d-amphetamine, see Figure 3. For reward, there was also a main effect of sex where males had higher reward delta power. In *mice*, there were no significant outcomes for either reward or punishment.

**Figure 3.**
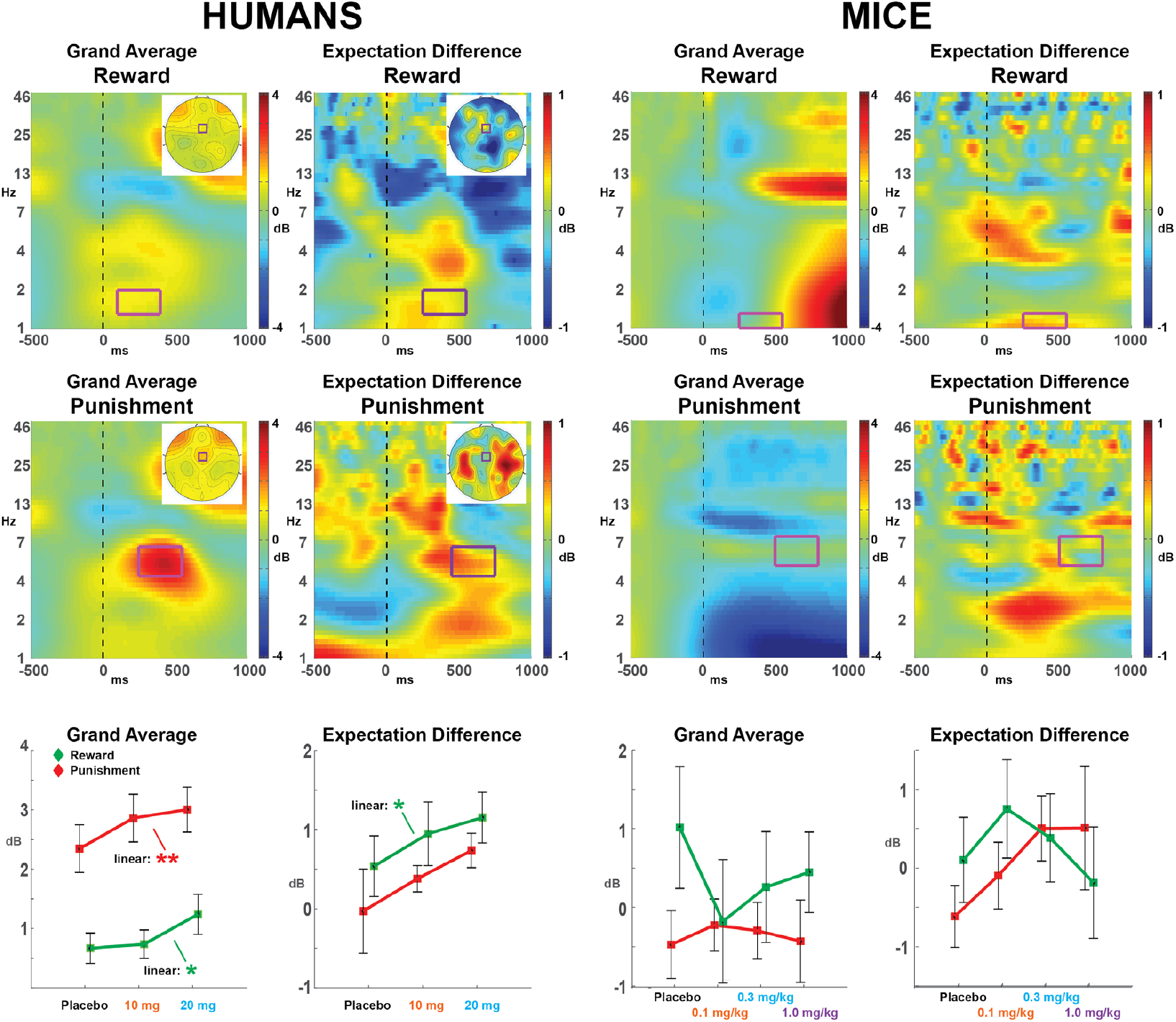
Time-frequency grand averages and expectation-related contrasts. Time-frequency plots of grand average and expectation difference (feedback from low vs. high probability conditions) at FCz in humans or the anterior lead in mice. The magenta box shows the tf-ROI. Human figures: grand average topographic plots: +/- 5 dB, expectation difference topographic plots: +/- 0.5 dB. Line plots (mean +/- SEM) detail the drug effects on each tf-ROI. *p<.05 **p<.01.

### TF-ROI: Expectation

All statistical results for tf-ROI expectation contrasts (low probability minus high probability) are reported in Table 1. In *humans*, there were main effects of drug with a linear trend for increased reward delta power with increasing dose of d-amphetamine. This effect was not present in the corresponding punishment condition. There was also a main effect of sex (higher power in males) and an interaction between drug and sex, where this expectation difference declined slightly in males and increased in females with increasing dose of d-amphetamine. In *mice*, there were no effects of d-amphetamine on tf-ROI expectation contrasts; however this is likely due to the poor performance learning the behavioral discriminations (Figure 1B). In sum, only humans displayed an effect of d-amphetamine on reward-related tf-ROI power.

### Small-scale replication of mouse ERP findings

We further aimed to replicate these appealing mouse ERP findings in a separate small-scale cohort with easier learning discriminations (a single pair of stimuli: target 80% correct vs. non-target 20% correct). Nine mice were run on placebo or 0.3 mg/kg of d-amphetamine across multiple sessions. All mice learned to perform around 80% accuracy with no difference between drug conditions. Supplemental Figure S4 shows that this drug-related enhancement of ERP amplitude could be replicated, although it did not achieve statistical significance (t(8)=1.64, p=.14, *d*=.41). However, the p-value is a poor metric for assessing replicability; effect sizes and confidence intervals are more useful for assessing the utility of an experimental outcome (Halsey et al. 2015; Colquhoun 2017). The effect size from this small-scale replication cohort is larger than the one for cohort described in the main test (i.e. Figure 2): a paired t-test between placebo and 0.3 mg/kg d-amphetamine on reward ERP amplitudes was t(27)=2.02, p=.05, *d*=.23.

## Discussion

This report validated our previous finding of a common electrophysiological marker of cortical reward processing in mice and humans as seen here during placebo (Cavanagh et al. 2021). The current work went further however, by demonstrating similar sensitivity of both species to increasing doses of d-amphetamine, demonstrating pharmacological predictive validity for this biomarker of reward sensitivity. While our previous cortical marker was based on the expectation-modulated spectral power, this study only found common cross-species amphetamine effects in the grand average ERP component. Unfortunately, the specific influence of d-amphetamine on learning-related spectral power in mice could not be definitively assessed due to poor learning. Yet, these findings already suggest that the influence of d-amphetamine is intrinsic to the generation of the RewP in both humans and mice.

As noted earlier, the RewP is sensitive to learning-related prediction errors, yet it is elicited by any rewarding feedback even in un-learnable environments (Foti and Hajcak 2009; Angus et al. 2015; Proudfit 2015; Mulligan and Hajcak 2017). This tendency appears to be preserved in the mice, who performed above chance but not at a level indicative of active successful learning. Our second small-scale replication cohort with good performing mice suggests that the enhancing effect of d-amphetamine on the RewP is reliable. However, two major issues remain to be addressed: 1) the specificity of d-amphetamine effects on reward vs. punishment conditions, and 2) the effect of d-amphetamine on reward signal generation vs. learning-related enhancements of this reward signal.

### Specificity vs. Generality of d-Amphetamine Effects on Reward

In humans, there were general facilitating effects of d-amphetamine, although the effects were strongest for the reward-related conditions. The reward-related ERP, grand average tf-ROI, and expectation tf-ROI were all significantly affected by d-amphetamine in humans, whereas punishment-related effects were smaller and had some different overall trends than the RewP. However, it is important to note that smaller effect sizes are not indicative of specific functional dissociation. In mice, there were also general facilitating effects of d-amphetamine on reward ERP amplitudes. Why did d-amphetamine affect spectral power in humans but not mice? One reason might be the much lower peak frequency of the mouse delta band phenomenon, which makes effective time-frequency quantification more difficult, particularly in the absence of a learning-related contrast (see section below). In sum, the cumulative cross-species similarities demonstrate that d-amphetamine boosts reward-related EEG signals in humans as well as mice, although the specificity of this effect to rewards could not be definitively determined.

Previous evidence of dopaminergic sensitivity of these cortical signals in humans has also been mixed, likely due to methodological reasons. Some studies have examined pharmacological effects on the punishment-related EEG signal or the difference between reward and punishment conditions, and have generally failed to find effects of the dopamine D2 receptor antagonists haloperidol (Forster et al. 2017) or sulpiride (Mueller et al. 2014a; Lueckel et al. 2018) unless moderated by genotype (Mueller et al. 2014b). However, conceptual issues about different generative systems underlying reward and punishment EEG signals casts doubt on the suitability of this common “difference wave” contrast (Meyer et al. 2017; Brown and Cavanagh 2020). One experiment revealed a reduction in the RewP in response to the dopamine D2/3 receptor agonist pramipexole (Santesso et al. 2009), yet there are no other studies of parametric or enhancing effects of dopaminergic agents specific to the RewP in humans. A recent report observed that although the RewP is specifically diminished in Parkinson’s disease, L-dopa administration did not alter this signal (Brown et al. 2020a). This surprising lack of acute dopaminergic influence might be due to combined issues of cortical degeneration in Parkinson’s and the differential influence of L-dopa on striatal vs. cortical dopamine tone (Cools 2006). The findings from this report extend the specificity of this extant literature in humans, demonstrating a clear influence of cortical dopaminergic agonism on the RewP and associated EEG signatures of reward receipt.

### Reward Signal Generation vs. Learning-Related Enhancements

Our previous investigation of cortical feedback signals was limited by the need to create a well-controlled analytic contrast within each species (i.e. low vs. high probability outcomes corresponding to high vs. low reinforcement prediction error), without interference from different sensory or imperative stimuli (Cavanagh et al. 2021). Since rewarded actions for mice were signaled with a tone indicating strawberry milkshake but punishments were signaled with the house lights, outcome valences were inherently incomparable. The expectation-defined tf-ROI was thus an excellent beginning to the identification of a common reward-specific biomarker; however, the expression of learning-related modulations is only a part of the relevant variance in this reward signal.

Expectation-related contrasts are ideal for identifying the specificity of pharmacological manipulations on these reward signals: the significant tf-ROI enhancement of delta power in humans due to d-amphetamine is powerful evidence for a selective and specific enhancing effect of dopamine agonism on this reward-responsive biomarker. Unfortunately, the absence of a learning effect in mice makes the null effect in this contrast less informative. Yet positive findings in the ERP component generation of this signal across both species is still particularly meaningful.

An emerging literature is revealing that appetitive motivation and affective value boost the RewP outside any influence of learning or expectation (Threadgill and Gable 2017; Brown and Cavanagh 2018; Peterburs et al. 2019; Huvermann et al. 2021; Pegg et al. 2021). This dissociation in affective vs. informational value might be critical for understanding altered reward dysfunction: for instance, major depression is associated with a diminished RewP but no change in the information representation of positive prediction error, suggesting that mood affects variance related to the generation of the signal and not the learning-dependent modulation (Cavanagh et al. 2018).

### Limitations and Future Directions

Cross-species comparisons in electrophysiological responses are complicated by a number of factors. There are differences between species in cortical homologies and the scale of neural activity in each recording type (i.e. scalp vs. dura). Still, increasing evidence suggests comparable midfrontal cortical structures (Balsters et al. 2020; Schaeffer et al. 2020) and confirms empirical similarities in EEG responses between humans and rodents (Narayanan et al. 2013; Ehlers et al. 2014, 2020; Warren et al. 2015; Featherstone et al. 2018; Robble et al. 2021). There are also inherent differences between species in task experience, motivation, and difficulty. We addressed these methodological difficulties with a highly constrained analytic strategy: we used the same task and the same time-frequency region of interest (tf-ROI) within our ‘learn-confirm’ strategy between experimental phases (Cavanagh et al. 2021).

While this methodological conservation was designed to constrain interpretation, continued methodological advancements will be required to optimally hone this field of research. For example, our small-scale replication study identified a better technique for enhancing learning (e.g. one 80/20 pair). Our findings reported here only contained a single electrode in humans with a single dura lead in mice. While this theory-driven reduction of spatial dimensionality was appropriate for the constrained methodological approach of this study, it offers only a fraction of assessable EEG activities in each species. In the future, depth recordings might provide more signal to noise than dura screws while simultaneously revealing the source generators of this reward-specific signal.

The use of pharmacological manipulation is a powerful tool for causal inference, although this technique also has inherent limitations. It is important to note that d-amphetamine enhances release of both norepinephrine and serotonin in addition to dopamine, and it is thus conceivable that some of the drug effects detected in this study reflect increases in non-dopaminergic neurotransmission. Yet in contrast to difficult and expensive human clinical trials, future rodent studies using this described paradigm could provide a relatively simple test of the aminergic specificity of the RewP.

### Summary

This study demonstrated that the RewP is a pharmacologically valid biomarker of reward sensitivity across species. Moreover, we provided a confirmatory test on the role of dopamine in the generation of the RewP. The RewP appears to be a trans-diagnostic biomarker of reward responsiveness, learning, and valuation. The cross-species translatability of this bio-signal will further bolster our mechanistic understanding of reward-related disfunctions in major depression, schizophrenia, and Parkinson’s disease.

## Data Availability

All data and Matlab code to re-create these analyses are available at OpenNeuro.org, accession #[WILL BE COMPLETED UPON ACCEPTANCE]

## Author Contributions (CRediT)

**JFC**: Conceptualization, Methodology, Software, Formal analysis, Writing - Original Draft, Funding acquisition

**SO**: Investigation

**JAT:** Investigation, Data curation, Supervision, Project Administration

**JEK:** Investigation, Data curation, Supervision, Project Administration

**BZR:** Investigation, Data curation, Supervision, Project Administration

**JAN:** Investigation, Data curation, Supervision, Project Administration

**JS:** Investigation, Data curation, Supervision, Project Administration

**DG**: Investigation

**SGB**: Conceptualization, Methodology, Investigation, Data curation, Supervision, Project Administration, Writing - Review & Editing, Funding acquisition

**GAL**: Conceptualization, Methodology, Resources, Writing - Review & Editing, Funding acquisition

**NRS**: Conceptualization, Methodology, Writing - Review & Editing, Investigation, Data curation, Supervision, Project Administration, Funding acquisition

**JWY**: Conceptualization, Methodology, Writing - Review & Editing, Funding acquisition

**JLB**: Conceptualization, Methodology, Resources, Writing - Review & Editing, Funding acquisition

## SUPPLEMENTAL MATERIALS

**Supplemental Table 1.** Characterization of the human cohort. WRAT: Wide Range Achievement Test; MCCB: MATRICS Comprehensive Cognitive Battery.

|  |  |
| --- | --- |
| Age (Mean(SD)) | 22 (4.82) |
| Race (%):<br>Caucasian<br>Asian<br>Pacific Islander/Native Hawaiian/Alaskan<br>Mixed race | 39<br>26<br>17<br>15 |
| Gender: M:F | 12:11 |
| Education (Mean (SD)) | 14.26 (1.74) |
| Smokers: non-smokers | 0:23 |
| WRAT score (Mean (SD)) | 108.13 (11.61) |
| Caffeine intake in mg/day (Mean (SD)) | 165 (231.6) |
| MCCB composite T-score (Mean (SD)) | 48.17 (9.06) |

**Supplemental Table 2.** Timeline for each experimental day for human participants.

| Time (hours) | Procedure |
| --- | --- |
| 0830 | Check-in; urine toxicology / pregnancy test; check for medication/lifestyle changes; vital signs & symptom rating scales; standardized breakfast |
| 0900 | Pill administration (placebo vs D-amp 10 mg vs. D-amp 20 mg) |
| 0930 | Vital signs & symptom rating scales |
| 1055 | Vital signs & symptom rating scales |
| 1100 | PLT with simultaneous EEG recording |
| 1215 | Vital signs & symptom rating scales |
| 1220 | Lunch |
| 1315 | Vital signs & symptom rating scales |
| 1415 | Vital signs & symptom rating scales |
| 1510 | Vital signs & symptom rating scales |
| 1610 | Vital signs & symptom rating scales |
| 1630 | Vital signs & symptom rating scales; checkout |

**Supplemental Table 3.** Mixed linear model outcomes of behavioral performance accuracy for humans and mice.

|  | Main:<br>Drug | Main:<br>Block | Main:<br>Sex | Drug*Block | Drug*Sex | Block*Sex | 3-Way |
| --- | --- | --- | --- | --- | --- | --- | --- |
| Human | F=3.12<br>df=55.12<br>p=.08 | **F=32.01**<br>**df=73.74**<br>**p<.001** | F=2.88<br>df=85.16<br>p=.09 | F=.76<br>df=94.85<br>p=.38 | **F=5.68**<br>**df=55.12**<br>**p=.02** | F=.74<br>df=73.74<br>p=.39 | F=1.2<br>df=94.85<br>p=.28 |
| Mouse | F=.44<br>df=141.83<br>p=.51 | F=.45<br>df=200.47<br>p=.50 | F=1.89<br>df=192.89<br>p=.17 | F=1.07<br>df=174.81<br>p=.30 | F=.12<br>df=141.83<br>p=.73 | F=.44<br>df=200.47<br>p=.51 | F=.02<br>df=174.81<br>p=.89 |

## Materials and Methods

### Animal subjects: pre-training & surgery

All operant behavior was conducted in a custom acrylic chamber measuring 21.6 × 17.8 × 12.7 cm housed within a sound- and light-attenuating box (Med Associates, St. Albans, VT). One end of the chamber contained a house-light, a tone generator, an ultra-sensitive lever, and a magazine with an attached peristaltic pump for delivering milk liquid reward (Nesquik (S.A., Vevey, Switzerland), non-fat powdered milk (Walmart INC., Bentonville, AR USA), and water). The other end of the chamber was fit with a touch-sensitive screen (Conclusive Solutions, Sawbridgeworth, UK) covered by a black acrylic aperture plate allowing two or two 2 × 5 cm touch areas separated by 0.5 cm and located at a height of 6.5 cm from the floor of the chamber. Stimulus presentation in the response windows and touches were controlled and recorded by the K-Limbic Software Package (Conclusive Solutions, Sawbridgeworth, UK).

Prior to training mouse body weights were slowly reduced and then maintained at 85% of their free-feeding body weight. Mice were acclimated to the testing room and the strawberry milk reward by providing 3 drops of reward per mouse on a weigh boat in the home cage for 15 min per day for 3 days. Mice were then habituated to the operant chamber and learned to collect liquid reward from the magazine wherein 30uL of strawberry milk was dispensed every 30 secs after each collection (measured by a head dip into the magazine). Mice that retrieved 10 rewards within 15 min were moved to bar training which required mice to press the lever for reward; criteria was 30 presses in 30 mins. Next mice moved to touch training, which required mice to initiate trials with a bar press and then respond to the presentation of a white (variously shaped) stimulus in 1 of the 2 windows on the touchscreen. The stimulus persisted until a response was made and touches to the blank window had no effect. Touching the stimulus resulted in a 1 second tone and reward delivery. Mice capable of initiating, touching, and retrieving 30 rewards in 30 mins moved to punish training. Punish training was identical to touch training except that touches to the blank window resulted in a 10 sec house light on time-out. Incorrect responses in this stage were followed by correction trials in which the same stimuli and left/right positioning was repeated until a correct response was made. Criteria was set at 75% correct responses in 30 mins.

Mice trained on the three-image set task in the main text next underwent surgery, while mice on the small-scale replication study (Fig. S4) learned to discriminate between two equi-luminescent images. Target images (Fan or Marble) had an 80% probability of reward delivery and a 20% probability of punishment. The non-target image was punished for 80% of trials and rewarded the other 20%. Image-reward contingencies were counterbalanced across mice. Once mice achieved 70% responses to the target image (regardless of rewards or punishments) over 2 days, they were single housed and given 3.0g of chow in preparation for stereotactic surgery.

During surgery, mice were anesthetized with 3% isoflurane and placed in a stereotaxic alignment system (Kopf Instruments, Tujunga, CA, USA) for surgical implantation of dura-resting skull screws (0.078” stainless steel machine screws (00.90-078-M-SS-PS US Micro Screw Seattle, WA, USA). Screws were placed targeting medial prefrontal cortex (mPFC: AP +2.80, ML +0.00) the primary motor cortex (AP +0.75, ML +1.50), and posterior parietal association cortex (PtA: AP −1.94, ML +1.50) with a cerebellar ground. Silver wire leads soldered to the pins of Omnetics connectors (Omnetics, MN, USA) were wrapped securely around each corresponding screw and secured to the skull using dental cement (Stoelting, IL, USA). After 4 days of recovery, body weight reduction resumed and mice were given post-surgery reminder sessions consisting of the last pre-training regimen to ensure retention of pre-training criterion. Once mice re-attained criteria of 70% target image selection they were moved to tether training which consisted of attaching the recording cable to the electrode cap under light isoflurane 15 mins prior to testing. Here mice learned to respond to stimuli with restricted movement and were required to maintain criteria of 70% target image selection before moving to recording. During each recording session, electrophysiological activity was captured via a multichannel acquisition processor (PlexControl, Plexon, Dallas, TX, USA) at a sample rate of 1000 Hz and behavior automatically video-tracked via headstage-integrated LED to exclude trial irrelevant behavior (CinePlex, Plexon, Dallas, TX, USA). Relevant task events and behaviors were time-coded as event markers on the recording file via TTL pulse from the behavioral software, and were confirmed via video tracking.

## Results

### Humans: Effect of d-amphetamine on autonomic activity and symptom ratings

D-amphetamine significantly increased heart rate from baseline with the maximum effect from 255 - 370 mins. Main effect of drug F_(2,44)_= 15.5, p<0.0001, time F_(6,132)_= 9.52, p<0.0001 and drug x time F_(^12, 264^)_ = 3.039, p=0.0005. Post-hoc significant 0-10 mg (p =0.0017) 0-20 mg (p<0.0001) and 10-20 mg (p=0.0351). D-amphetamine significantly increased diastolic blood pressure from baseline with the maximum effect from 115 - 430 mins. Main effect of drug F_(2,44)_= 4.33, p=0.02, time F_(6,132)_= 8.5, p<0.0001 and drug x time F_(12,264)_ = 2.126, p=0.02. Post-hoc significant 0-20 mg (p=0.0052). D-amphetamine significantly increased systolic blood pressure from baseline with the maximum effect at 115 min and then 255 - 430 mins. Main effect of drug F_(2,44)_= 11.22, p=0.0001, time F_(6,132)_= 5, p=0.0001 and drug x time F_(^12, 264^)_ = 2.328, p=0.007. Post-hoc significant 0-10 mg (p =0.04) 0-20 mg (p<0.0001) and 10-20 mg (p=0.012). There was no effect of d-amphetamine on symptom ratings of anxiety (F<1), happiness (F<1) or drowsiness (F_(2,42_)=1.38,ns), though ratings immediately after EEG testing revealed a significant reduction in drowsiness with d-amphetamine (Main effect of drug F_(2,42)=_3.91, p<0.03; 0 vs. 10 mg: p<0.009). When asked to identify what condition they were in (drug or placebo), participants were 83% correct on placebo days, 65% correct on 10 mg days, and 74% correct on 20 mg days.

### Mice: Replication of Simple Expectation Effect

In order to quantitatively inform the emerging literature on this novel bio-signal, we aimed to directly replicate the statistical test from our previous study (Cavanagh et al. 2021). We collapsed the placebo, 0.1 mg/kg, and 0.3 mg/kg conditions and ran a t-test on the tf-ROI delta band power between expectation conditions. Even in the absence of statistically robust learning across the cohort, this contrast was t(27)=1.43, p=.17, *d*=.12, the mean dB difference was .41, and the correlation between this expectation difference dB and aggregate accuracy for each mouse was rho(27)=.16, p=.43. These values are similar to the two cohorts in our foundational paper (*d*’s= .60, .18; dB differences =.65, .65; cohort 1 correlation rho=.38), providing evidence of reproducibility of this expectation-dependent delta-band reward feature.

**Figure S1.**
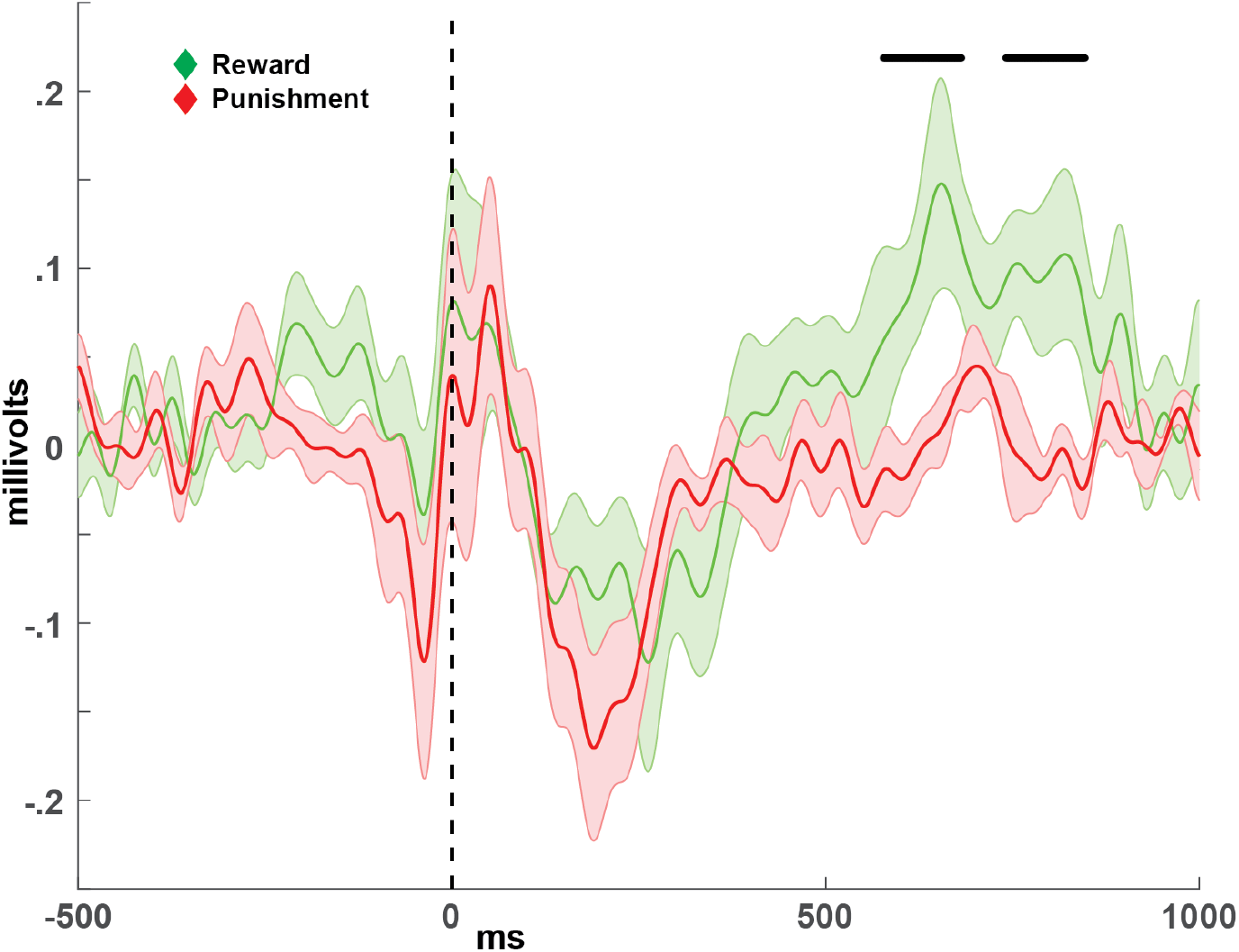
Feedback-locked ERPs from mice in our previous investigation (Cavanagh et al. 2021). Black bars indicate statistically significant differences between conditions (N=12 mice, p<.05, uncorrected for multiple comparisons). The large voltage deflection around time 0 are motion artifacts. Similar to human studies (see discussion), the reward-specific activity differs in time between cohorts. Here the reward-specific activity peaks 500-800ms post-reward.

**Figure S2.**
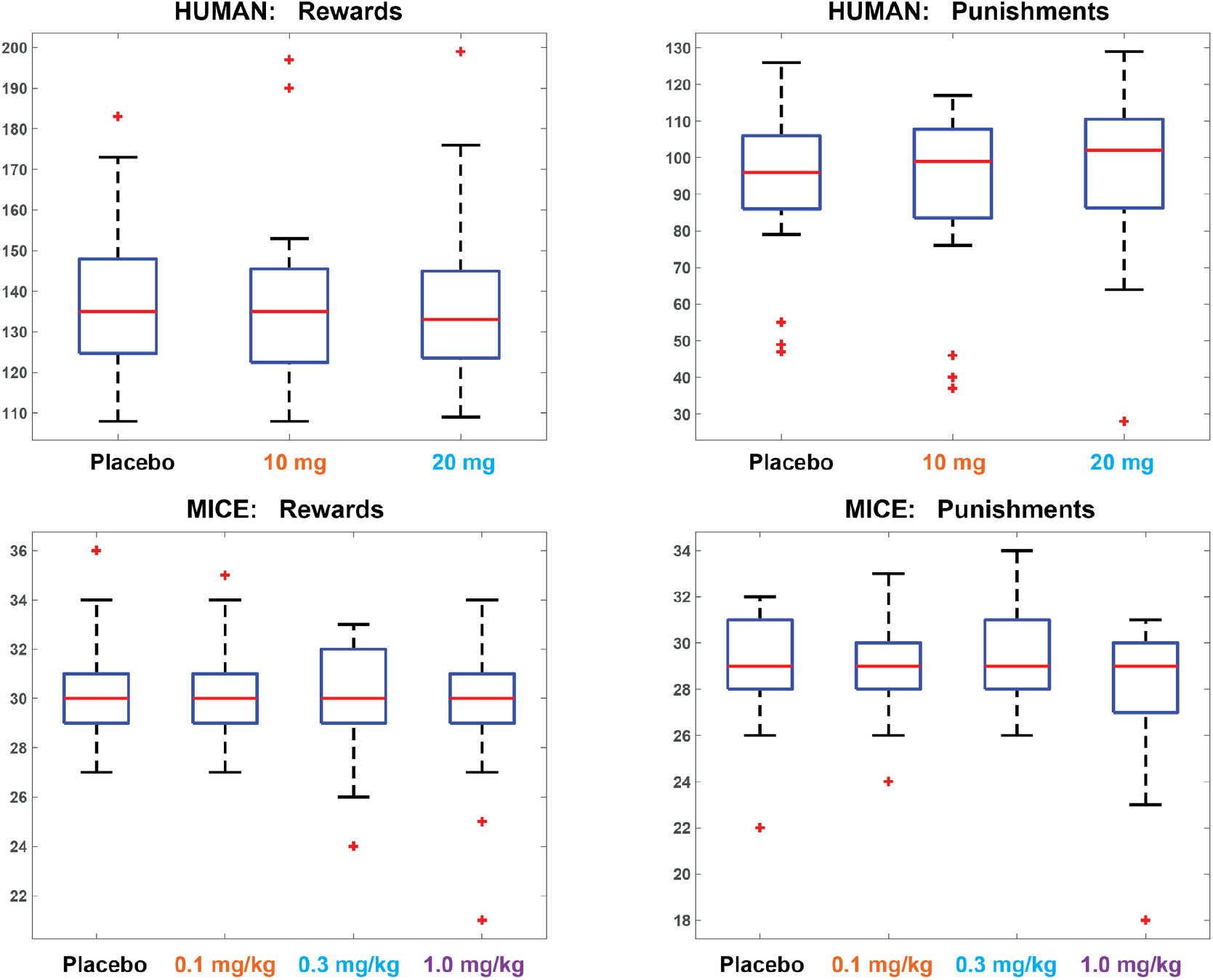
Number of epochs for each condition.

**Figure S3.**
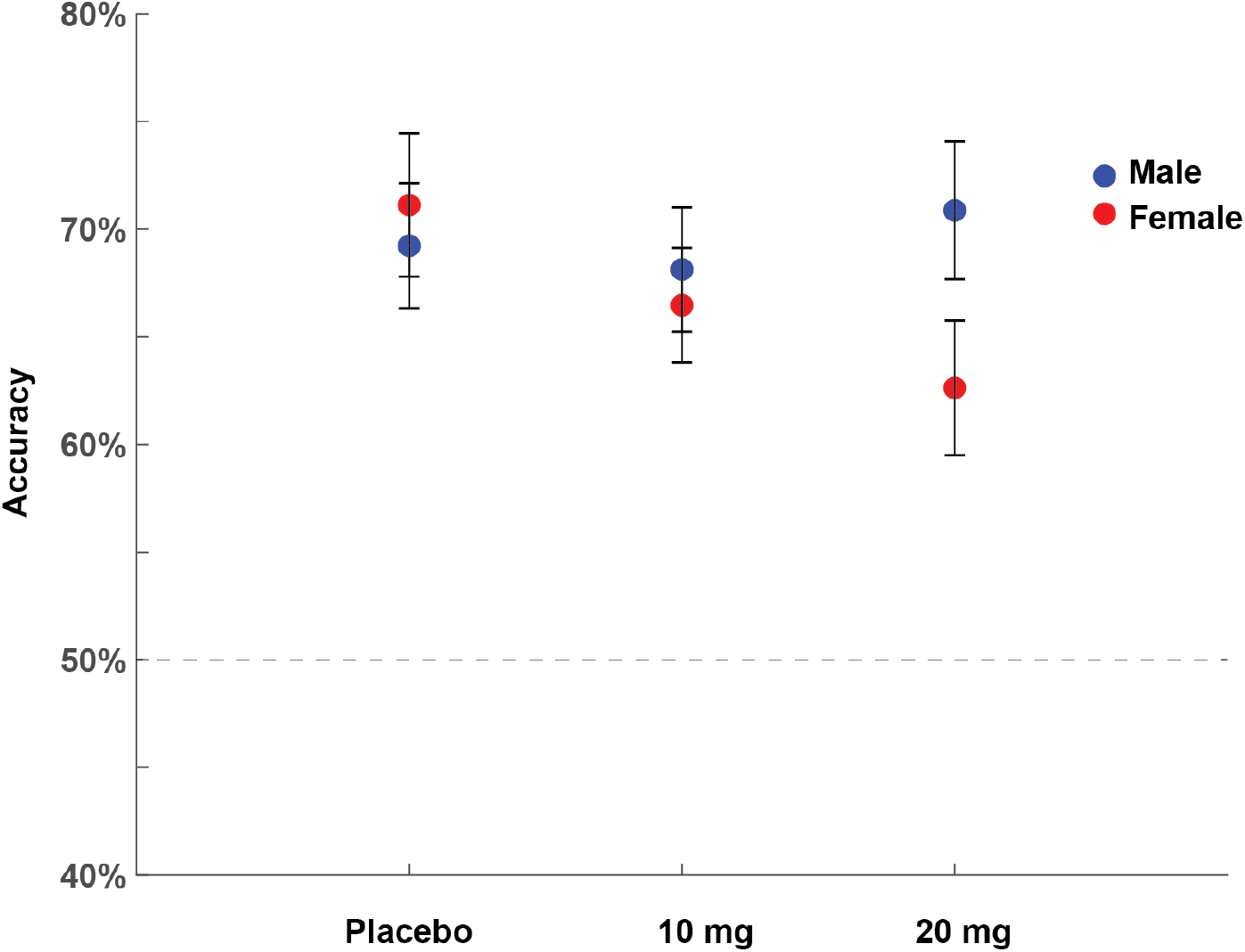
Human behavioral performance collapsed across block. This figure shows the sex*drug interaction whereby increasing dose of d-amphetamine (X-axis) had no effect in males but caused decreased performance accuracy in females.

**Figure S4.**
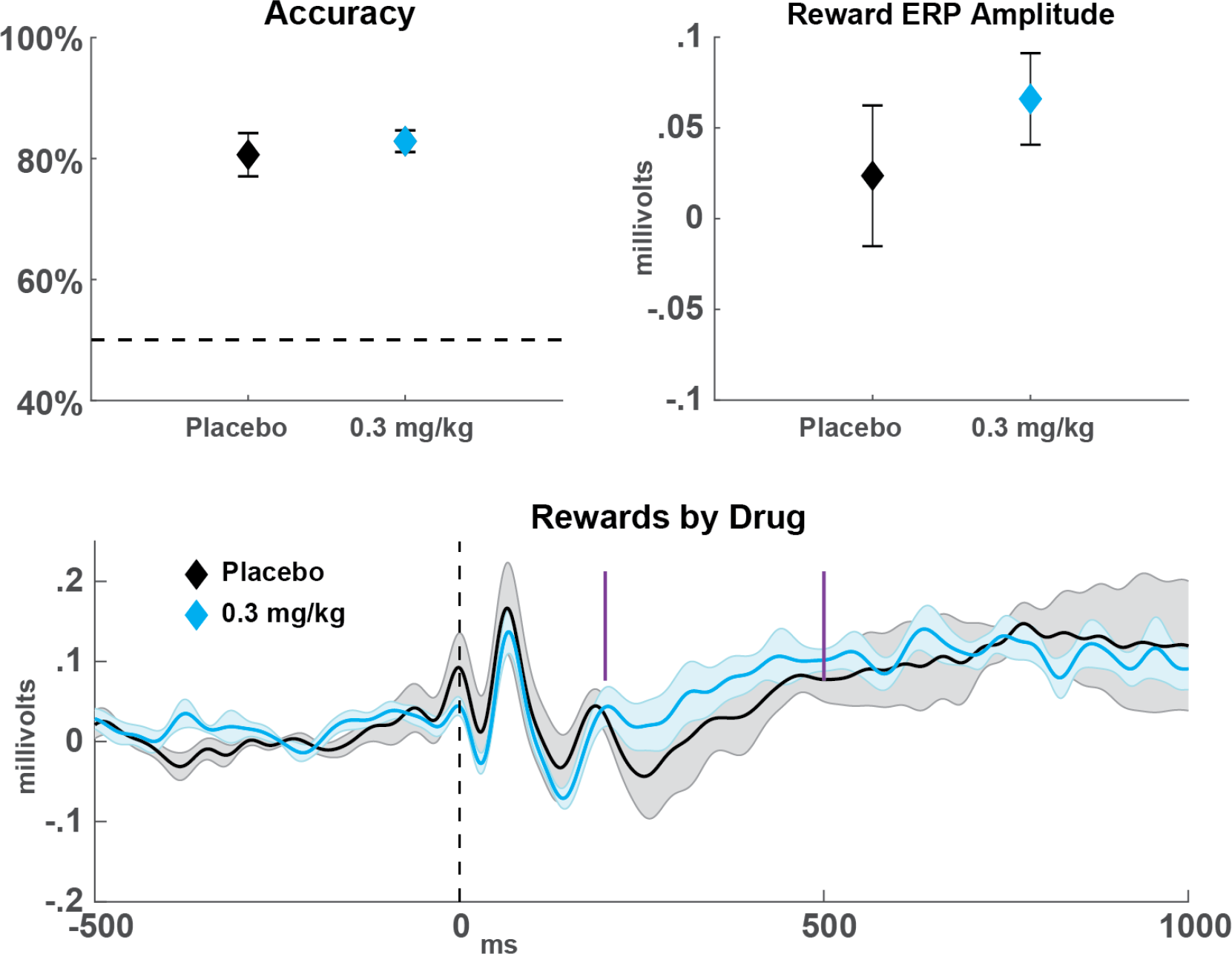
Small-scale replication study. An additional cohort of mice was run with a simpler reward structure and only two drug conditions (placebo, 0.3 mg/kg). All mice were all able to learn to discriminate a single pair of stimuli (target: 80% correct / non-target: 20% correct) at approximate matching levels, with no difference between drug sessions (t(8)=1.03, p=.33). Of these nine mice, six had eight sessions each (four placebo), two had four sessions each (two placebo), and one died after two sessions (one placebo). The reward-specific component peaked earlier than both our prior investigation (Figure S1: 500:800 ms) and the current main text report (Figure 2: 400:600 ms); here the RewP was quantified at 200:500 ms. This analysis showed a trend in the expected direction, but this difference was not statistically significant (t(8)=1.64, p=.14, *d*=.41). However, this contrast has a larger effect size than the paired contrast for these two conditions from the main text: t(27)=2.02, p=.05, *d*=.23. In sum, these findings suggest that these ERP and amphetamine effects are reliable, but like in human ERP findings there is substantial inter-individual variability that affects component identification and statistical power. All plots are mean +/- SEM.

